## supplemental information for "Normative age modelling of cortical thickness in autistic males"

### Site and participant information

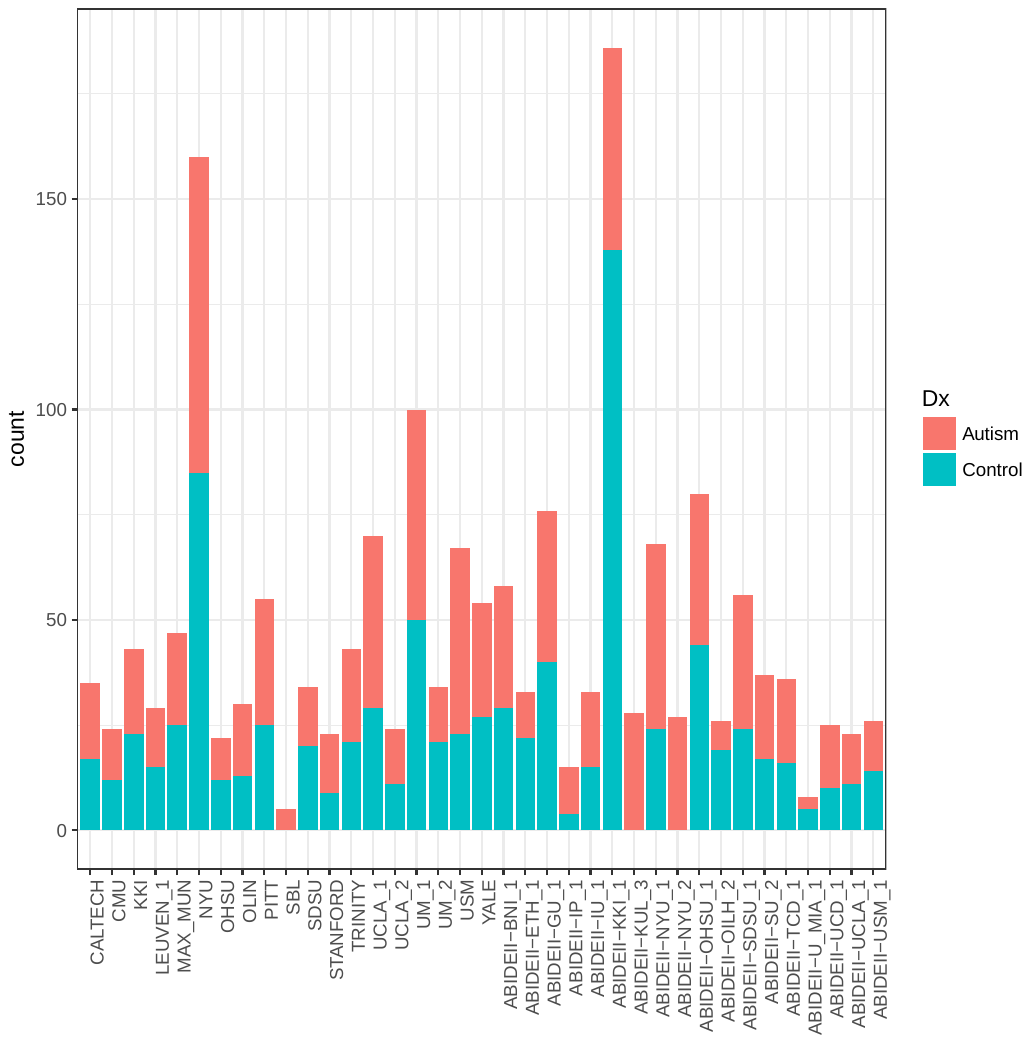

##### **Supplemental Figure S1:** Site distribution

### Quality control

To assess quality of Freesurfer reconstructions we computed the Euler index (Rosen et al., 2018). The Euler number is a quantitative index of segmentation quality and has shown high overlap with manual quality control labelling. In the full sample we found a small but significant difference in both hemispheres (Figure S2) with the autism group having overall worse scan quality (d = 0.176 and d = 0.187 for left and right hemisphere respectively). We excluded all subjects with a Euler score of 300 or higher in either hemisphere. After thresholding based on Euler indices and after re-running the sample consisted of 870 individuals with autism and 870 neurotypical individual, matched on Age and IQ.

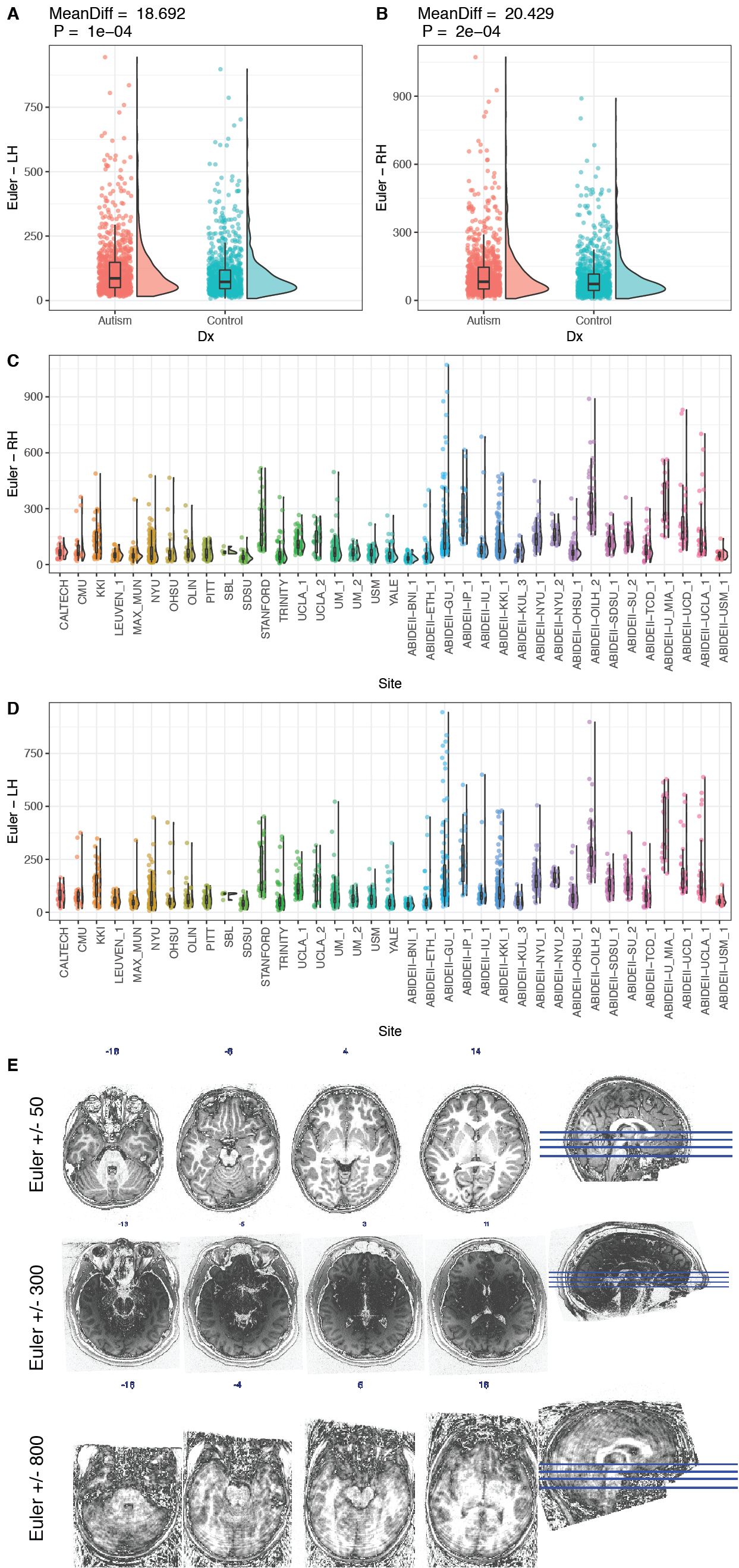

##### **Supplemental Figure S2:** Euler quality control

Panels A and B list the Euler indices for both left and right hemisphere, p-values for group differences were established with two-sided permutation testing (20000 permutations), Cohen’s d was computed using custom R code <https://github.com/mvlombardo/utils/blob/master/cohens_d.R>. Panels C and D show the left and right hemisphere Euler distribution across the different sites included in ABIDE. Panel E shows 3 example scans for Euler indices of 50, 300 and 800 respectively. Raincloud plots were created using: <https://micahallen.org/2018/03/15/introducing-raincloud-plots/>

### Bootstrapping

To assess the reliability of this w-score we bootstrapped the normative sample (1000 bootstraps, with replacement) and computed 1000 bootstrapped w-scores for each individual and each brain region. To subsequently quantify the reliability of the w-score we computed an FDR corrected analogous p-value for each subject by computing the absolute position of the real w-score in the distribution of bootstrapped w-scores. The rationale being that if a real w-score would be in the top 5% of the bootstrapped distribution it would likely not be a reliable score (e.g. the score would be influenced by only a small subset of the normative data). The median number of brain regions per subject with a significant p-value was 1 (out of 308), indicating that the w-score provides a robust measure of atypicality.

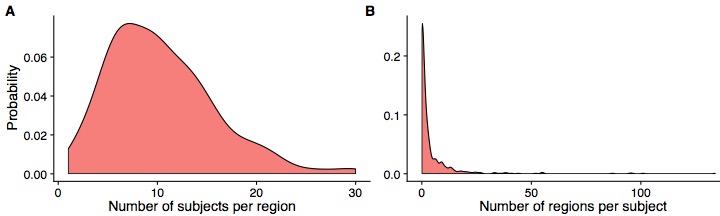

##### **Supplemental Figure S3:** bootstrap validation

Panel A shows the probability density distribution for the number of subjects likely to have an unreliable w-score in a given region. For a given region there is a median of 10 subjects (out of 714) for which the w-score is not reliable (e.g. has a p-value <0.05). Panel B shows the probability density distribution for the number of regions likely to have an unreliable w-score in a given subject. The median number of 'unreliable' brain regions per subject is 1.

### Multivariate clustering

Using t-Distributed Stochastic Neighbour Embedding (tSNE) was used to construct a distance matrix from the 308 ROI features. K-medoid clustering on this distance matrix was used to cluster subjects into maximally independent groups using the optimum average silhouette width (Hennig and Liao, 2013) to determine the optimal number of cluster. This identified an optimal number of two clusters. Finally, we re-ran k-medoid clustering with 2 clusters and explored the overlap these clusters gave with diagnosis by visualizing the clustering onto the 2-dimensional embedded space obtained from tSNE.

Despite the limited main diagnosis effect on CT over the majority of brain regions and the fact that only a small subset of individuals appears to contribute to this difference, it may still be possible that the multivariate patterning in CT may capture some diagnostic effect. Thus, we performed exploratory clustering analysis to determine if raw CT values across the whole brain could be used to delineate the ASD group from the TD group. In addition, we reasoned a data-driven clustering approach might also reveal subgroups within each group (Lombardo et al., 2016). Results from clustering the neighbour embedded raw CT scores are shown in Figure 6. As can be observed in panel B, the within-group heterogeneity is entirely captured by normative heterogeneity and the overall density plots for both groups are close to identical. The pattern we find when projecting the whole brain raw cortical thickness into a 2-dimensional embedding most closely resembles the 3^rd^ scenario outlined by Marquand and colleagues (Marquand et al., 2016), whereby disease related variation is nested with the normal variation. Our results show that, when it comes to whole-brain cortical thickness, the condition related variation is entirely nested within the neurotypical variation. Obviously, the present clustering and embedding approaches only provide one way of clustering or segregating case-control variation in cortical thickness. Other multivariate approaches that took into account a multitude of variables did reveal that multivariate clustering has the potential to identify subgroups (Hong et al., 2017). Additionally, other measures than CT might provide a different picture. In is interesting however to note that both dimensions were correlated with age. No correlations were observed with any of the other common phenotypic measures. Thus, this 2-dimensional embedding likely captures the variability in cortical thickness expansion and thinning over the course of development, but is not sensitive enough to pick up potential alterations in the overall trajectory of this process between groups.

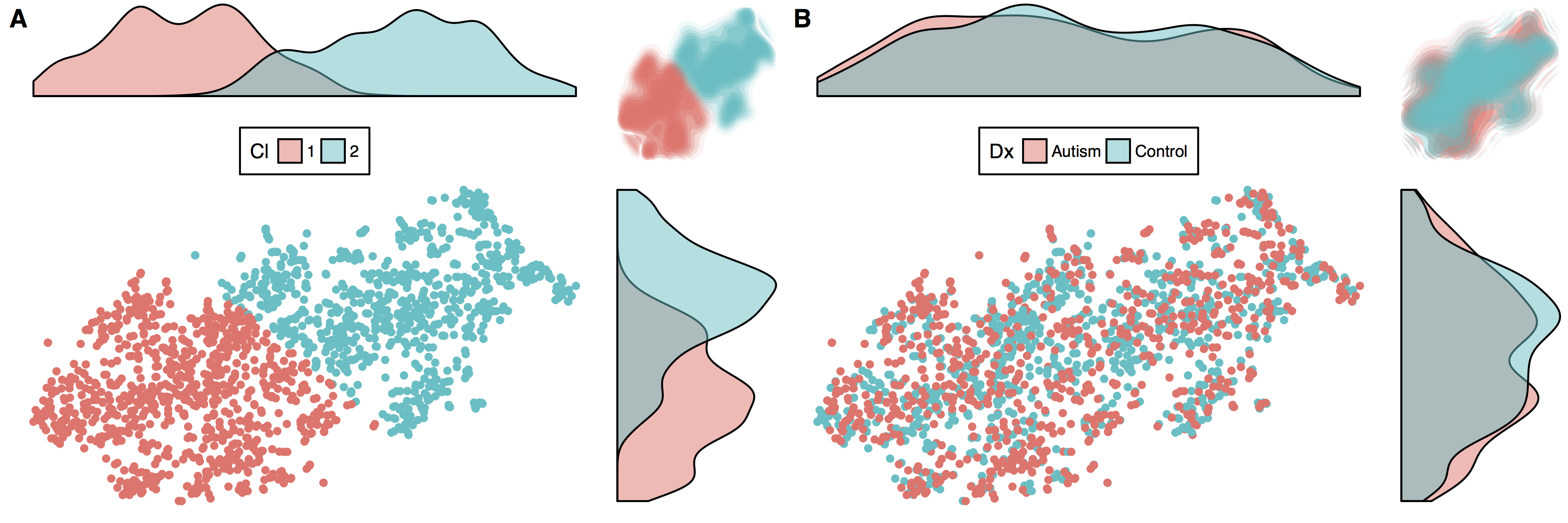

##### **Supplemental Figure S4:** tSNE Clustering

Panel A shows the results from k-medoid clustering of the 2D embedding of the raw CT values as achieved by tSNE. K-medoid clustering clearly identifies 2 clusters. However, as Panel B shows, these clusters did not provide any meaningful distinction on diagnosis.

### Individual summary ratios

To isolate subsets of individuals with significant age-related CT deviance, we used a cut-off score of 2 standard deviations (i.e. w >= 2 or w<=2). This cut-off allows us to isolate specific ASD patients with markedly abnormal CT relative to age-norms for each individual brain region. We then calculated sample prevalence (percentage of all ASD patients with atypical w-scores), in order to describe how frequent such individuals are in the ASD population and for each brain region individually. A sample prevalence map can then be computed to show the frequency of these patients across each brain region. We also wanted to assess how many patients have markedly atypical w-scores (beyond 2SD) across a majority of brain regions. This was achieved by computing an individual global w-score ratio as follows:

$$gW=\frac{\sum\left| \left( w \right) \right|>2}{\sum\left| \left( w \right) \right|<2}$$

We also computed global w-score ratios for positive and negative w regions separately.

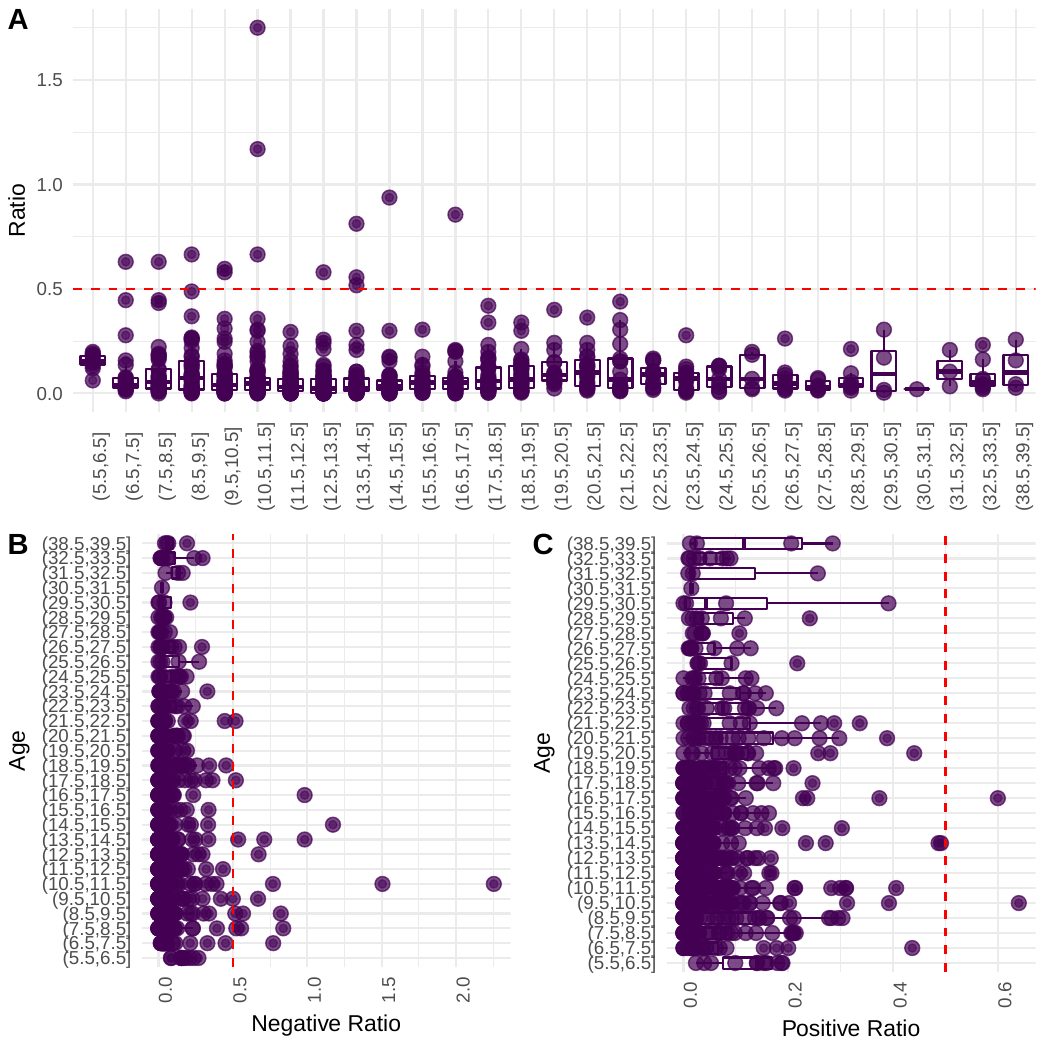

##### **Supplemental Figure S5:** Global individual W-Score ratios

Panel A shows the distribution of absolute global ratio scores for each age-bin. There is a total of 14 subjects for which the ratio score exceeds 0.5 meaning they have more atypical than typical regions. Panels B and C show the same but stratified for positive and negative outliers.

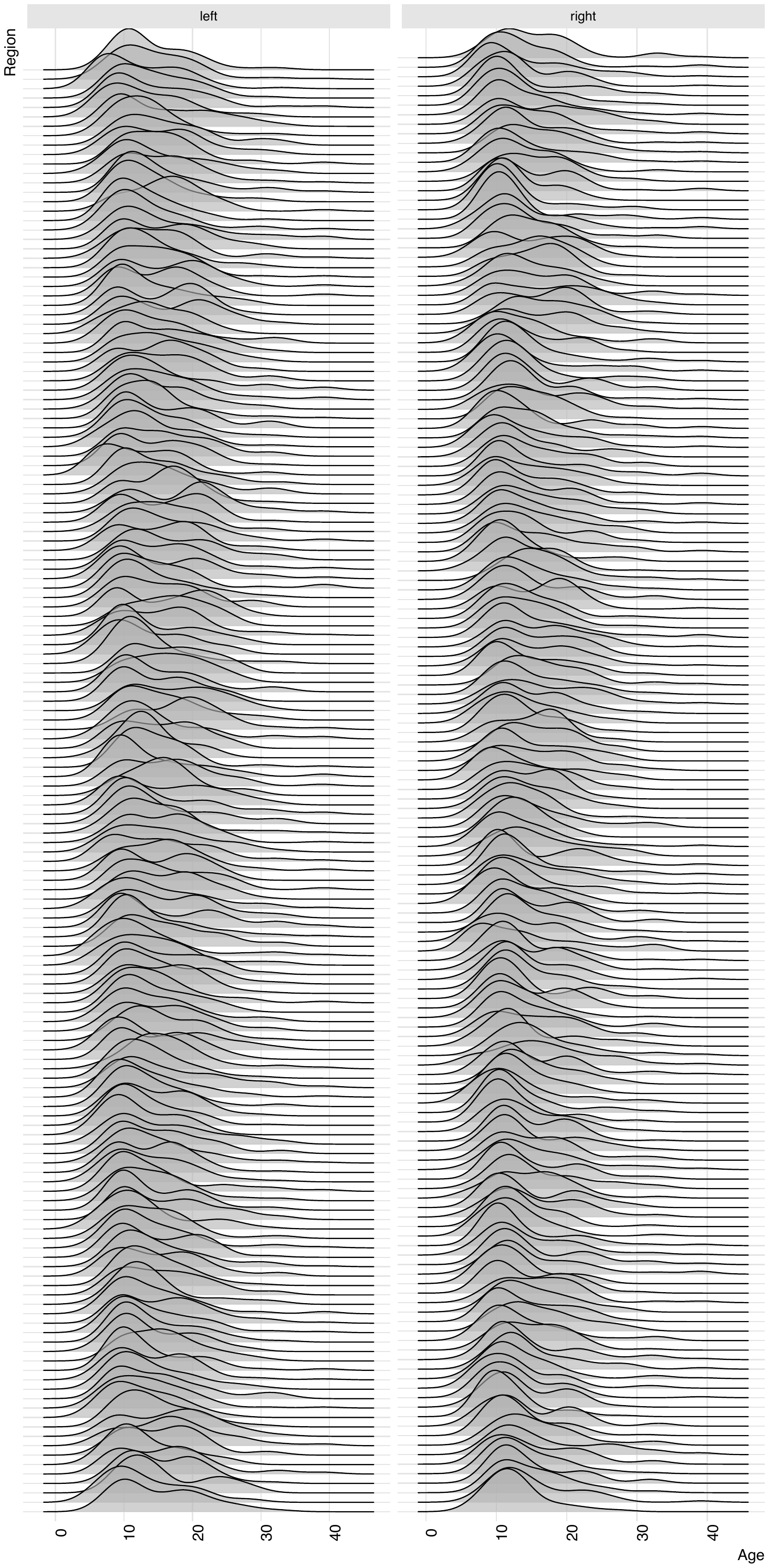

##### **Supplemental Figure S6:** Outlier age distribution per brain region

Probability density plots of the age of all outliers for each brain region. Left and right refer to left and right hemisphere.

### Comparison to centile modelling of normative deviation

In order to assess the sensitivity of our approach in the present data we implemented the aforementioned bootstrapping procedure to identify robustness of outlier detection. In addition, we also conducted a centiles estimation that is relatively standard in for example epidemiology (Visser et al., 2009), similar to quantile rank maps (Chen et al., 2015) and arguably less sensitive to small sample uncertainty. Both approaches showed high significant correlation in determining whole-brain w-score ratios (r=0.87, p=4e-119 and r=0.66, p=5.7e-39 for ABIDE I and ABIDE II respectively).

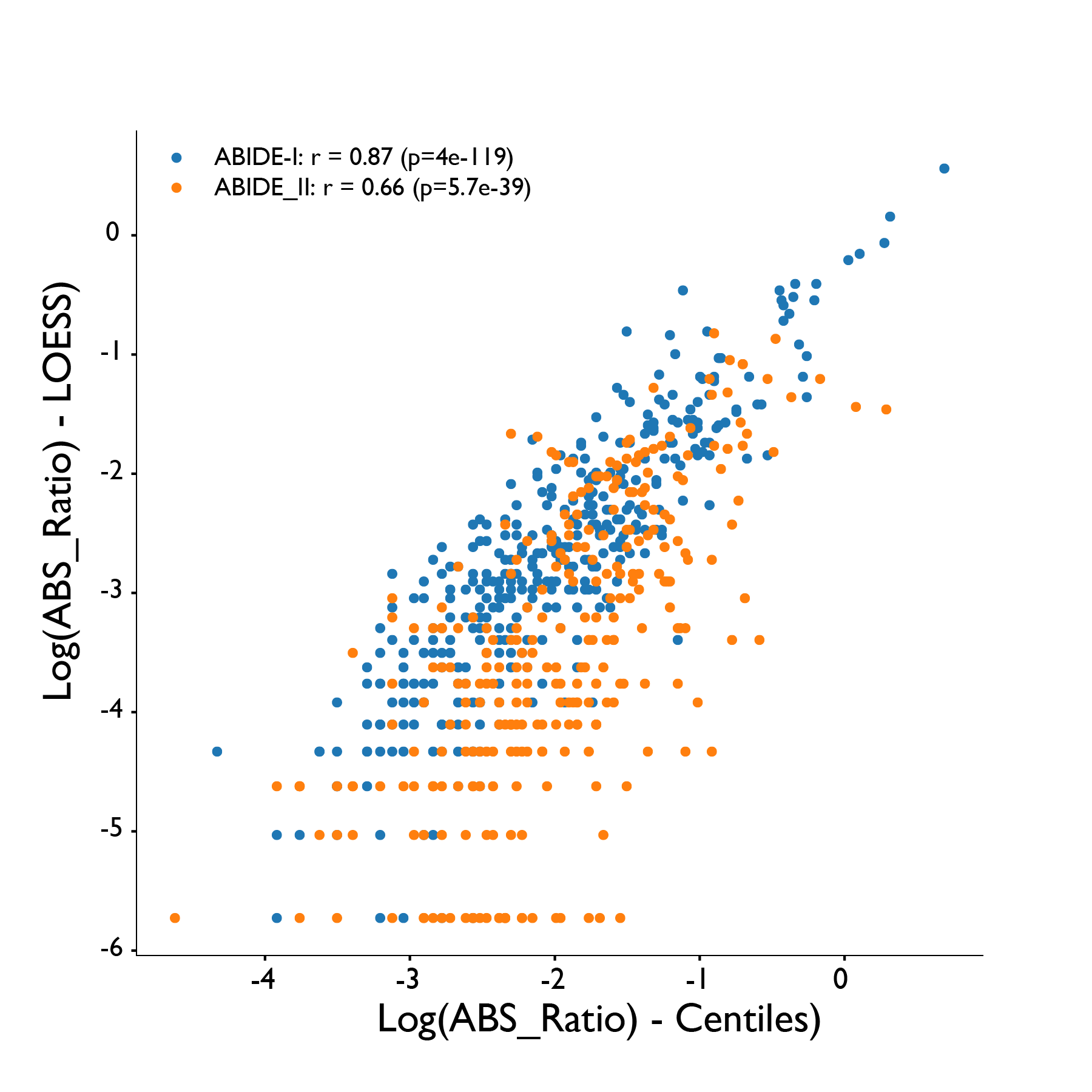

##### **Supplemental Figure S7:** Centile vs LOESS regression

Scatterplot of the absolute ration of atypical regions in both ABIDE I and ABIDE II as computed using LOESS regression and centiles estimation.

### Age-related CT deviance relationships with SRS and ADOS

An additional advantage of the use of normative modelling over the traditional case-control modelling is that we can use the individualized deviation as a novel metric for finding associations with phenotypic features. Here we used w-scores to compute Spearman correlations for the most commonly shared phenotypic features in the ABIDE dataset: ADOS, SRS, SCQ, AQ, FIQ and Age. After correcting for multiple comparisons across phenotype and region (6 phenotypic measures * 308 regions = 1848 tests) we identified a number of brain regions that survive multiple comparison corrections for the SRS and ADOS scores (supplementary figure S8). SRS is associated with w-scores primarily in areas of lateral frontal and parietal cortex, while ADOS is associated with w-scores primarily in lateral and inferior temporal cortex. Notably, these regions are largely different from regions that appear to show on-average differentiation in case-control and w-score analyses.

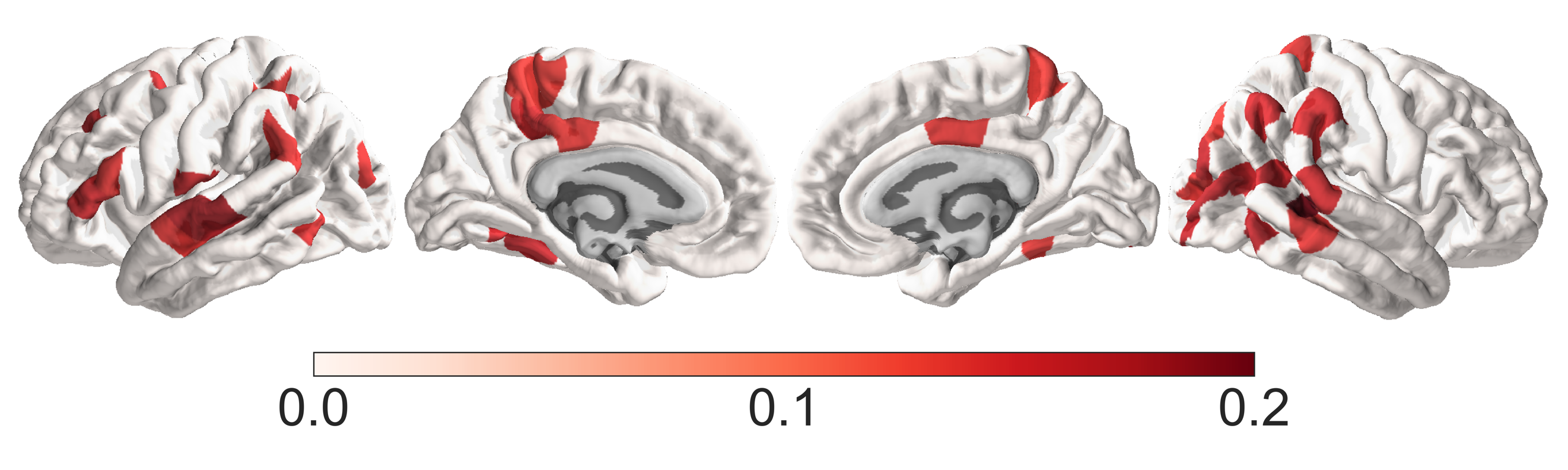

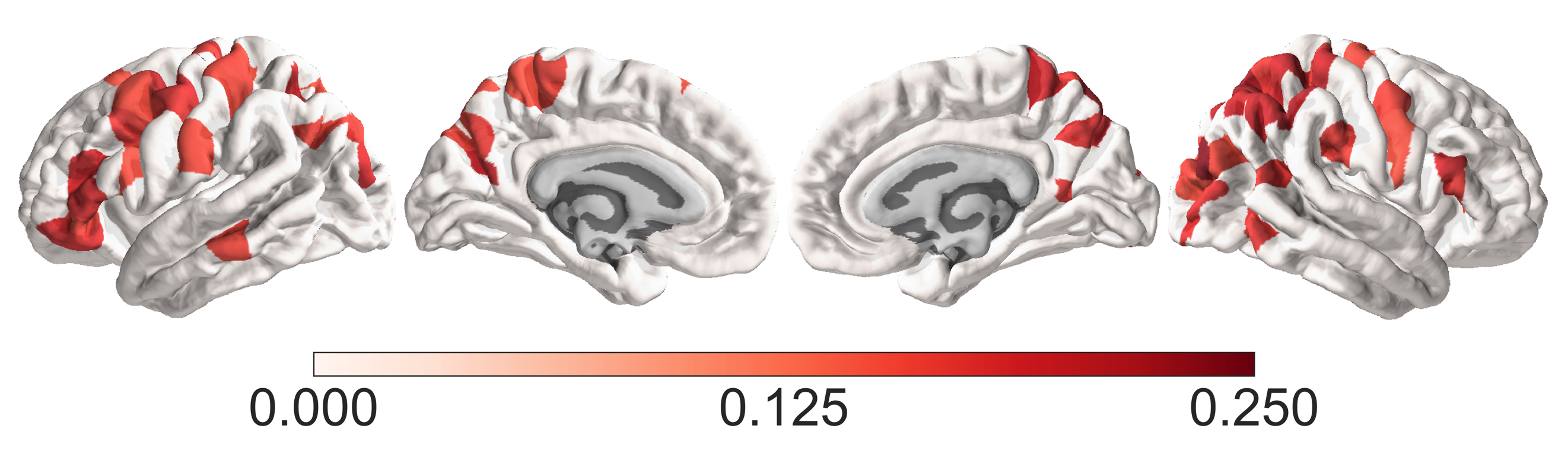

##### **Supplemental Figure S8:** Phenotype – W-Score correlations

Spearman correlations between ADOS and w-score in the top panel. The lower panel shows the same for the SRS.

### Surface area, LGI and Volume

We applied the same approach to quantify outlier contribution and assess overall variance contribution in surface area, LGI and cortical volume. All three metrics consistently showed strong influence of sex and scanner site as important covariates (S8). In addition, cortical volume also showed a strong influence of age. For all three measures all canonical case-control differences, derived from standard LME modelling, disappear when region-wise outliers were removed (S9-S11). This strongly suggest that the majority of broad case-control differences were driven by a subgroup of individual outliers.

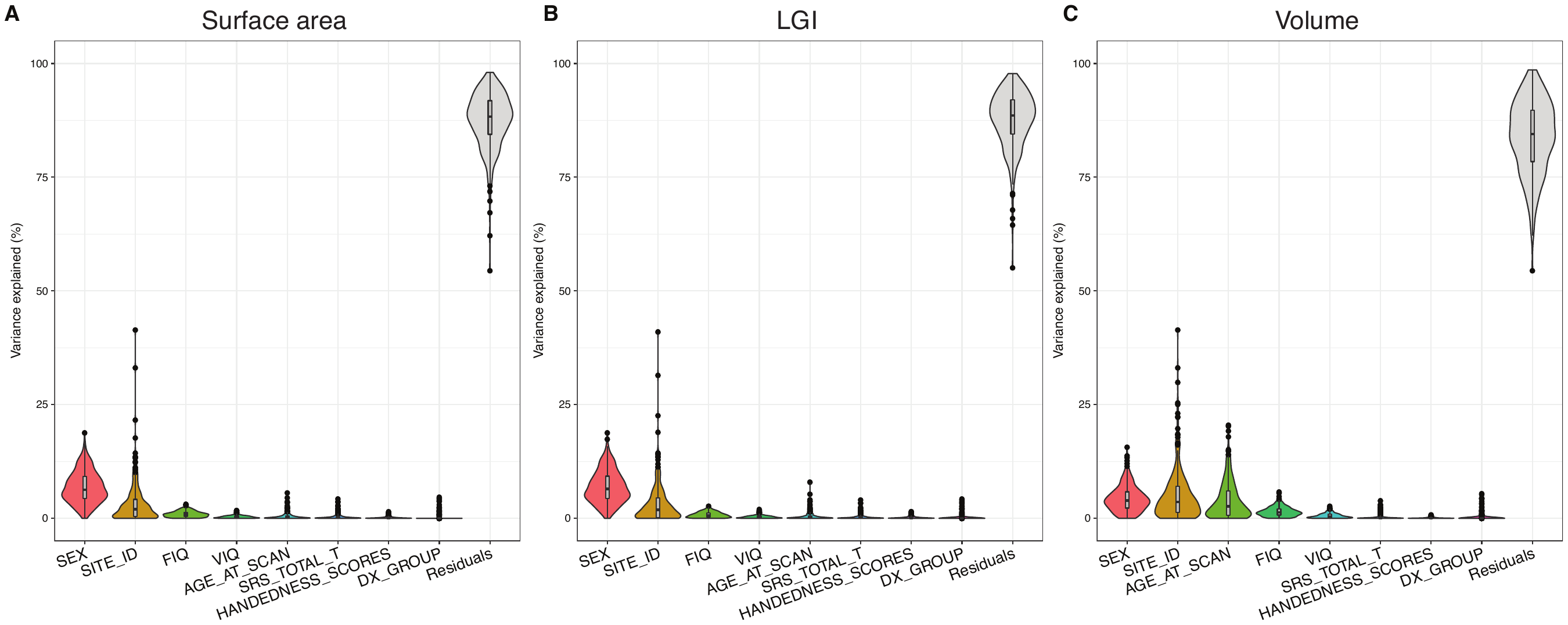

##### **Supplemental Figure S9:** Variance contribution across measures

Gender and scanning site are the dominant sources of covariance

#### Volume

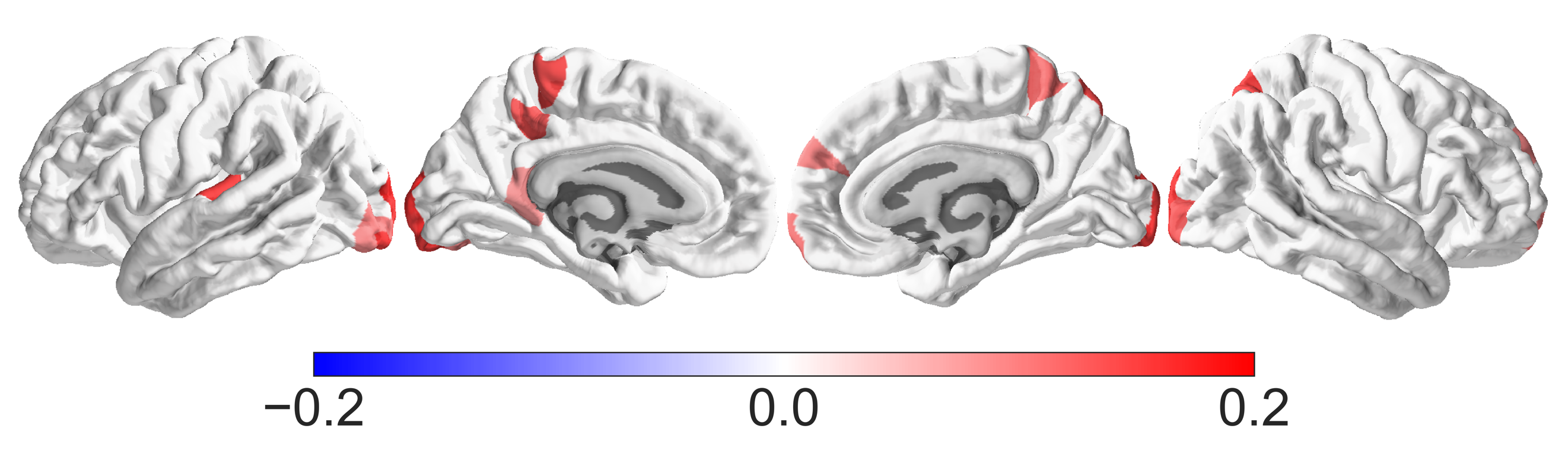

##### **Supplemental Figure S10:** Canonical case-control comparisons for cortical volume

The top panel shows the canonical case-control output. No regions passed FDR when region-wise outliers were removed nor on the one sample w-score test.

#### LGI

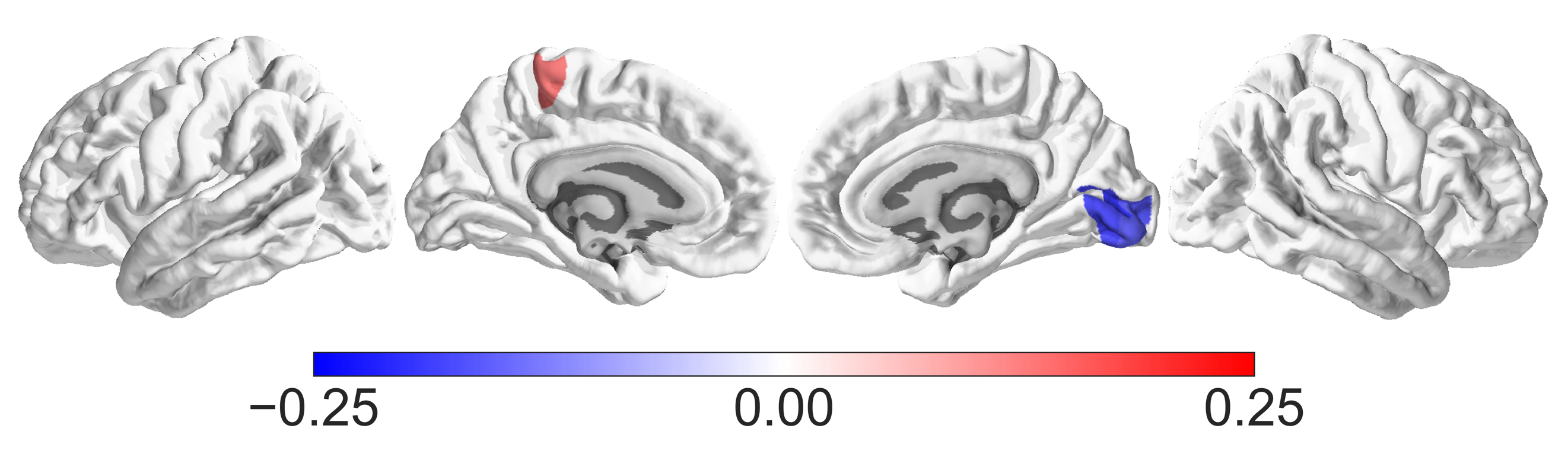

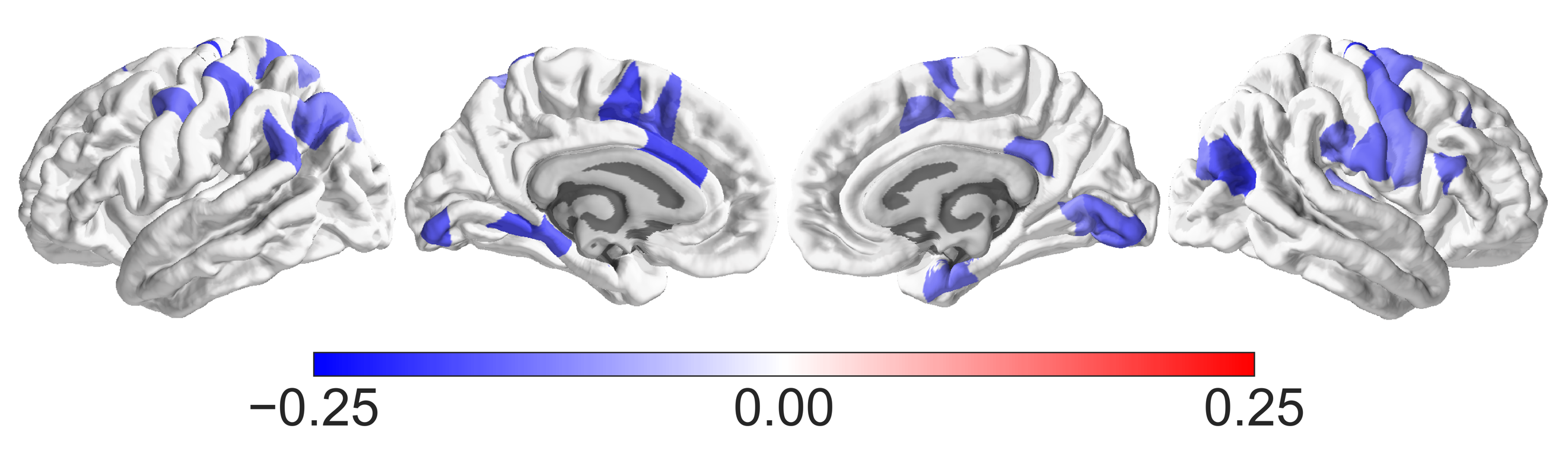

##### **Supplemental Figure S11:** Canonical case-control comparisons for local gyrification

The top panel shows the canonical case-control output. No regions pass FDR when region-wise outliers are removed however in the w-score one sample test (lower panel) there are a number of regions that show significantly smaller LGI in ASD.

#### Area

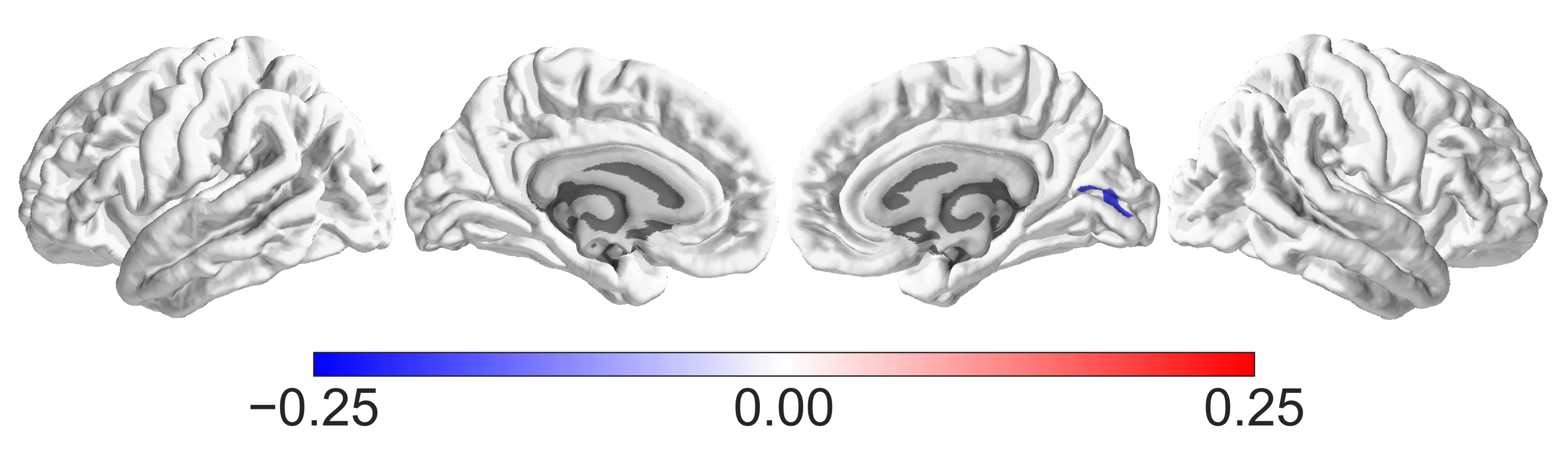

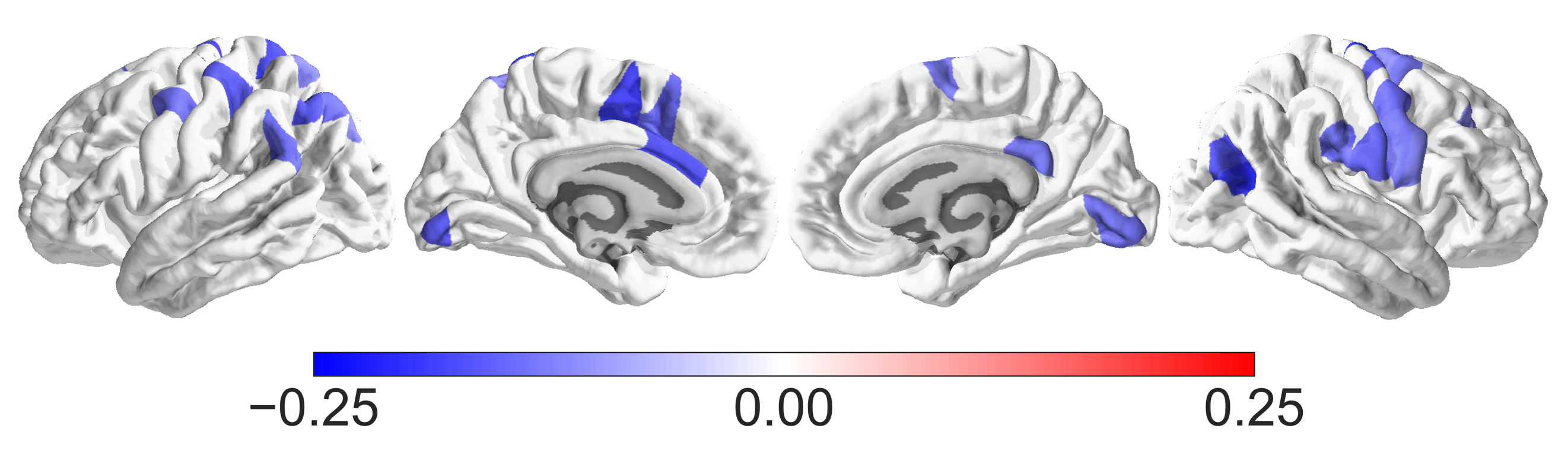

##### **Supplemental Figure S12:** Canonical case-control comparisons for surface area

The top panel shows the canonical case-control output. No regions pass FDR when region-wise outliers are removed however in the w-score one sample test (lower panel) there are a number of regions that show significantly smaller surface area in ASD.
